# Biofilm rupture by laser-induced stress waves increases with loading amplitude, independent of location

**DOI:** 10.1101/827964

**Authors:** Kaitlyn L. Kearns, James D. Boyd, Martha E. Grady

## Abstract

Integral to the production of safe and biocompatible medical devices is to determine the interfacial properties that affect or control strong biofilm adhesion. The laser spallation technique has recently emerged as an advantageous method to quantify biofilm adhesion across candidate biomedical surfaces. However, there is a possibility that membrane tension is a factor that contributes to the stress required to separate biofilm and substrate. In that case, the stress amplitude, controlled by laser fluence, that initiates biofilm rupture would vary systematically with location on the biofilm. Film rupture, also known as spallation, occurs when film material is ejected during stress wave loading. In order to determine effects of membrane tension, we present a protocol that measures spall size with increasing laser fluence (variable fluence) and with respect to distance from the biofilm centroid (iso-fluence). *Streptococcus mutans* biofilms on titanium substrates serves as our model system. A total of 185 biofilm loading locations are analyzed in this study. We demonstrate that biofilm spall size increases monotonically with laser fluence and apply our procedure to failure of non-biological films. In iso-fluence experiments, no correlation is found between biofilm spall size and loading location, thus providing evidence that membrane tension does not play a dominant role in biofilm adhesion measurements. We recommend our procedure as a straightforward method to determine membrane effects in the measurement of adhesion of biological films on substrate surfaces via the laser spallation technique.

**Graphical Abstract:** 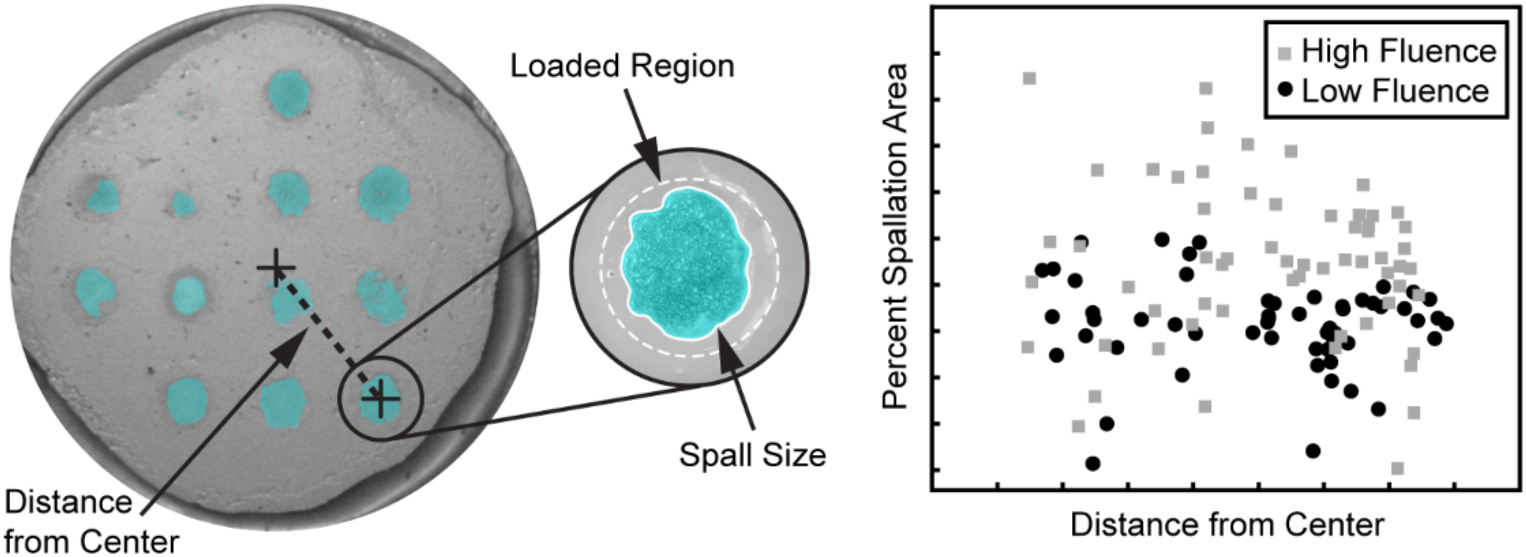

## 1. Introduction

Biofilms are thin coatings of bacteria that are encased in a gel-like material called extracellular polymeric substances, or EPS [1, 2]. The EPS is excreted from bacteria, and its mechanical properties are affected by many factors including nutrients in the environment, shear forces, and temperature [3–5]. The EPS contributes to the high adhesion strength of biofilms [6–8], which allows bacteria to thrive in a multitude of environments and often causes detrimental effects on the surfaces they inhabit. For example, bacterial biofilm formation on medical implants increases a patient’s risk of serious infection [9]. Among these devices are hip and knee implants [10], dental implants [11–13], artificial pacemakers [14], and fluid shunts [15], as well as non-permanent devices such as intravenous catheters [16], ventilators [17], and contact lenses and cases [18, 19]. By preventing the initial adhesion of biofilms to medical devices, the chance of associated bacterial infections will decrease dramatically [9]. Therefore, the study of biofilm adhesion to medical device substrates greatly benefits the medical community.

Several experimental methods have been developed to determine biofilm adhesion including counting methods [2, 20], atomic force microscopy [6], shear flow [21], and others [22, 23]. Key disadvantages of macroscale techniques such as shear flow are that the biofilm can respond to the force applied before adhesion can be measured and that multiple locations along the biofilm-substrate interface cannot be probed. The laser spallation technique was recently adapted to overcome these disadvantages by rapidly applying (within nanoseconds) a high amplitude stress wave, at multiple locations, on a single biofilm-substrate sample [24]. The laser spallation process begins with a substrate coated with a film for which an adhesion measurement is desired – in this case, a biofilm coating on a medical device surface. A single laser pulse impinges upon the uncoated side of the substrate, where the light energy is absorbed, confined, and transferred as a compressive stress wave that propagates toward the film coated side of the substrate. Once the stress wave arrives at the free surface, it reflects toward the interface in tension. If the magnitude of the tensile stress is greater than the adhesion strength of the coating-substrate interface, a portion of the coating is ejected in a process called spallation. The hole that remains in the coating after the film is ejected is referred to as the spallation region. The spallation region varies based on the laser loading fluence (energy per unit area), and the coating composition or growth conditions. For example, several authors [25–29] report a larger spall size at higher fluences when compared with the same coating loaded at lower fluences, though no systematic study has been presented. It has also been qualitatively shown that a loaded film with low adhesion strength would result in a larger spall size than that of a high adhesion strength film loaded at the same fluence.

Laser spallation has been implemented to measure adhesion of various coatings including metallic [25, 30–33] and polymeric [27, 29, 34] films, by obtaining the stress that causes film failure. The use of laser spallation on biological films such as cells and bacteria is a relatively newer field [35–37], thus these methods must be carefully examined to determine the validity of the measurements. Biofilms have the potential to exhibit membrane tension, a phenomenon that could influence adhesion measurements depending on the location at which a measurement is recorded. Membrane tension arises from the theory that a biofilm experiences a homeostatic tension that changes in amplitude with respect to distance from an edge. For example, on a membrane such as a drum, tension is higher at the edges than in the center. Because techniques like shear flow preclude probing multiple areas of the same biofilm, the influence of membrane tension on macroscale adhesion measurements has not been possible [14, 38]. The laser spallation technique enables the measurement of multiple locations across the entire biofilm and can determine the influence of membrane tension on adhesion measurements.

In this study, we provide the first systematic study of spallation region size with respect to location for biofilms on substrates. *Streptococcus mutans* is chosen as the model biofilm due to its promotion of other pathogenic bacteria and its presence in failing dental implants [39] and titanium is chosen because titanium and its alloys are the most widely used materials in structural implants [40–43]. The methods described in this paper can be employed by other investigators to deconvolute location bias due to membrane tension within biological films or edge effects for synthetic films during laser spallation testing.

## 2. Materials and methods

### 2.1 Substrate assembly

Glass slides (3”×1”×1.1 mm) coated one side with titanium (100 nm) and the opposite side with aluminum (300 nm) (Deposition Research Lab, Inc.) are cut into 3 approximately square pieces (1”×1”). The substrate is cleaned with a solution of 70% methanol in water and lens paper. Each square of substrate is placed on a spincoater (Specialty Coating Systems: Spincoat G3P-8) and the aluminum side is coated with approximately 2 mL of aqueous sodium silicate, or waterglass (Fisher Scientific). The spincoater ramps to 3000 RPM over 5 seconds, dwells for 40 seconds, and ramps to 0 RPM over 10 seconds, resulting in a waterglass thickness of 5 μm. The coated substrate is attached with silicone sealant (Dowsil 732) to the bottom of a petri dish (diameter 35 mm) to completely seal a hole (diameter 25 mm), which is cut into the bottom of the dish. The titanium-coated side faces upward in the dish so that the biofilm is grown on the titanium surface. The adhesive dries for at least 12 hours before adding media to the dishes. An example of a completed substrate assembly is shown in **Fig. 1a** and a substrate assembly with biofilm growth is shown in **Fig. 1b**.

**Fig. 1.**
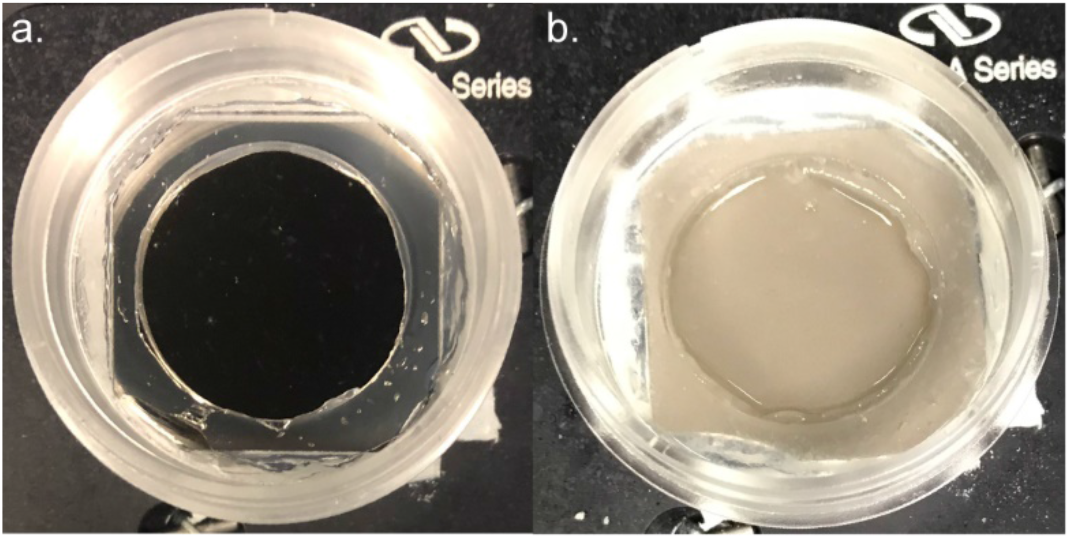
Photographs of (a) complete substrate assembly and (b) assembly with biofilm grown on the titanium surface. Both dishes are 35 mm in diameter with a hole of 25 mm in diameter.

### 2.2 Biofilm growth

THY broth (5 mL) is inoculated with frozen *S. mutans* stock in a 15 mL centrifuge tube. The inoculation is cultured in a water bath at 37°C for 24 hours. The mixture is diluted with additional THY, then vortex-mixed to dislodge bacteria attached to the bottom of the centrifuge tube. The final mixture has an optical density of 0.7 measured by a Thermo Scientific Genesys 30 Visible Spectrophotometer. In a separate 15 mL centrifuge tube, 0.75 mL of a 2 M sucrose solution is added to 15 mL of THY. Each substrate assembly is filled with 1 mL of *S. mutans* inoculum and 3 mL of the sucrose and THY mixture. The final concentration of sucrose in each dish is 75 mM. Sucrose aids in formation of EPS and the sucrose solution chosen for this study yielded the strongest adhesion rate when compared to four other sucrose concentrations [24]. The inoculated dishes are placed in an incubator at (37°C) for 24 hours. After a day of growth, biofilms form on the titanium surface and the remaining liquid is gently aspirated to avoid disturbing the biofilm. To measure thickness, the biofilms are dyed with Syto9 (Thermo Fisher Scientific) and imaged using a Zeiss LSM 880 upright multiphoton microscope. The z stack is processed using a commercially available biofilm analysis plugin (IMaris). Biofilm thickness is approximately 21.5 μm with a standard deviation of 2.3 μm.

### 2.3. Loading via laser-induced stress waves

The laser spallation setup consists of an Nd:YAG pulsed laser (λ = 1064 nm), a variable attenuator, a focusing lens, and an angled mirror (**Fig. 2**). The Nd:YAG emits a single pulse, which passes through the variable attenuator (Newport VA-BB series) to adjust the energy of each pulse. The pulse diameter is controlled by a focusing lens and the pulse is reflected upwards by a 45° angled mirror. Each substrate assembly is placed on a level sample holder that is oriented so that the pulse hits the bottom of the substrate, the waterglass layer, first. The waterglass acts as a confining layer, which amplifies the stress wave generated by impingement upon the aluminum layer [33]. The compressive stress wave is reflected as a tensile wave through the substrate and biofilm and, if the stress wave is of sufficient magnitude, causes the biofilm layer to spall or eject from the titanium surface. The spall size is dependent on both the growth medium and the final loading fluence (energy per unit area), but the maximum spall size is equal to the pulse spot size, which is set at a diameter of 2.2 mm for these experiments. Each biofilm is loaded multiple times in a 4 mm spaced grid pattern across the area of the biofilm by adjustment of micrometer-controlled translation stages (Thor Labs). Two distinct studies are performed: an iso-fluence study, in which each biofilm is loaded at a consistent fluence, and a variable fluence study, in which each biofilm is loaded multiple times at different fluences. The fluence values used to load the biofilms for the iso-fluence study are 55.6 mJ/mm^2^ and 79.4 mJ/mm^2^. In previous studies on *S. mutans* biofilms, both fluence values resulted in consistent rates of spallation, which allows facile spall size measurement [24]. For the variable fluence study, 6 biofilms are loaded at fluence values of 15.9 mJ/mm^2^, 23.8 mJ/mm^2^, 31.8 mJ/mm^2^, 39.7 mJ/mm^2^, 55.6 mJ/mm^2^, and 79.4 mJ/mm^2^ resulting in a total of 69 loading locations. By testing over a large range of fluences, we are able to capture the onset of spallation.

**Fig. 2.**
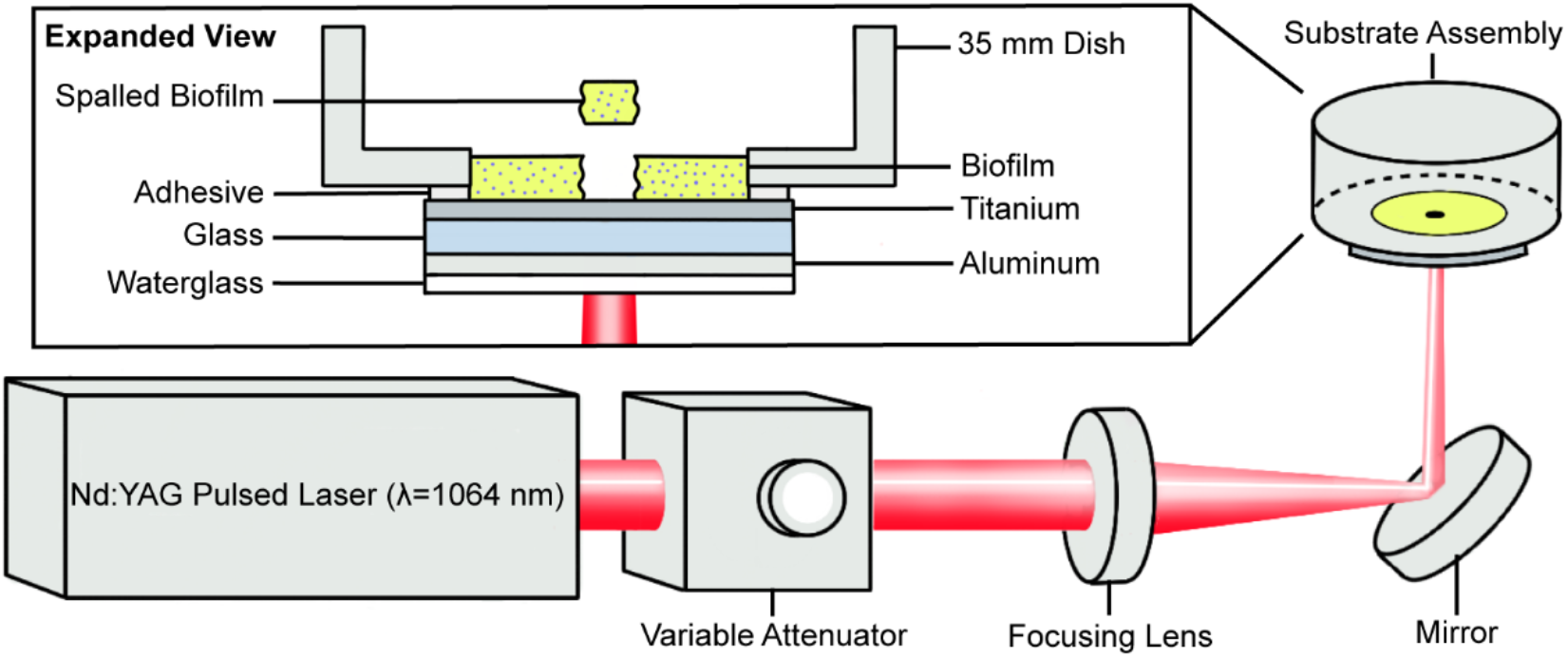
Schematic of components in the laser spallation setup. An expanded view of the substrate assembly is shown.

### 2.4. Spall size measurement

After loading, each spallation region is imaged with an Olympus SZ61 stereo microscope with an LC micro camera attachment, then analyzed in ImageJ [44]. In ImageJ, each image is converted from color to grayscale (**Fig. 3ab**), then a threshold is applied to locate the darkest pixels in the image (**Fig. 3c**). The image colors are inverted by converting to binary, and all the black pixels are selected (**Fig. 3d**), and the area of pixels is measured. The area is then converted into the percentage of possible spallation area, and that value is recorded. The image analysis process is shown in **Fig. 3**. and is performed for all 185 biofilm spallation regions included in this study.

**Fig. 3.**
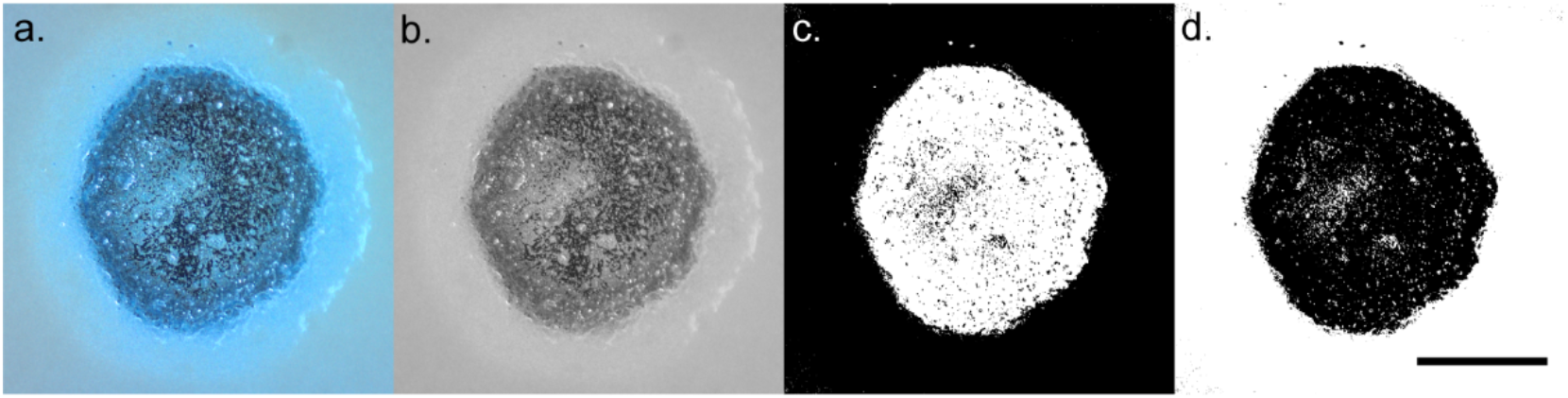
Process of spall size measurement on a region loaded at a fluence of 79.4 mJ/mm^2^: (a) raw optical image, (b) image is converted to 8-bit, (c) threshold is applied to locate darkest pixels, (d) image is converted to binary, and area of dark pixels is measured. The area measured is 2.81 mm^2^, which corresponds to 73% of the possible spallation area. Scale bar is 1 mm.

A significant number of laser spallation studies incorporate images of film failure by stress wave loading [24–29]. The sequence of images indicates an increase in spall size with each increment in laser fluence, however, this trend has not been previously quantified. To demonstrate applicability of our approach to analyze spallation images for synthetic films, we apply our procedure to failure of several films provided to us from two prior spallation studies: gold films transfer printed onto functionalized silicon substrates (55 loaded regions) [25], sol-gel films fabricated with a Pt/Ti superlayer to impart higher tensile stresses to cause spallation (4 loaded regions) [27]. The image analysis protocol outlined above is performed with the donated images from previous studies and the trend of spall size increase is presented alongside results from this biofilm study.

### 2.5. Distance from spallation region to biofilm centroid

Biofilms loaded at iso-fluence are analyzed to compare the resultant spall areas as a function of loading location. An image of the entire biofilm region within the field of view is obtained with a digital microscope camera (DinoLite Edge) with DinoCapture software. The biofilm is allowed to partially dry before the full image is taken to reduce glare. An example of a loaded and partially dried biofilm is shown in **Fig. 4**. In ImageJ, the circular outline is traced using a circle tool, and the center coordinates of the circle are recorded. Next, the coordinates of each spallation region are recorded. The distance from each spallation region to the center of the circle is calculated and recorded. **Fig. 4** consists of a biofilm with 13 loaded regions, four of which are shaded in cyan. The spall size for each region is ① 0.01 mm^2^ ② 2.08 mm^2^ ③ 2.08 mm^2^ and ④ 2.09 mm^2^. The cyan circle around region ① represents the maximum spallation region: a 2.2 mm diameter circle. A very small spall size is measured at region ①, which results from the small number of black pixels measured within the loading region. The cyan cross at the center of each spallation region marks the centroid of each region. The distance between each region (cyan cross) and the centroid (yellow cross) is ① 7.15 mm ② 6.77 mm ③ 6.12 mm and ④ 6.56 mm. This procedure is repeated across 5 biofilms at a fluence of 55.6 mJ/mm^2^ and 5 biofilms at a fluence of 79.4 mJ/mm^2^ for a total of 116 loaded regions.

**Fig. 4.**
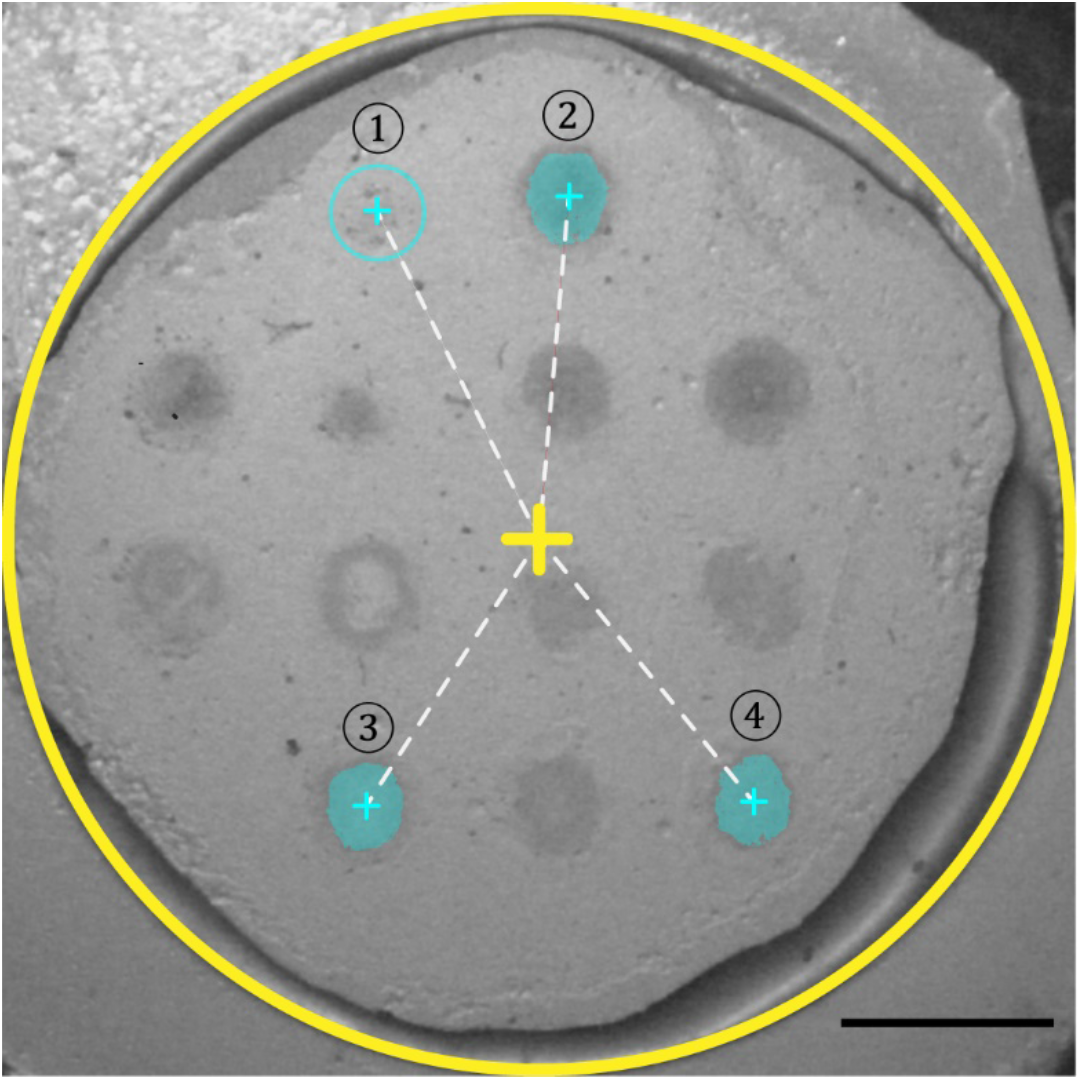
Optical image of a biofilm with 13 spallation regions loaded at a fluence of 79.4 mJ/mm^2^. The circular opening has a diameter of 25 mm, outlined in yellow with a cross at the center. Cyan regions correspond to example spallation regions, and dashed lines correspond to the distance between each spallation region and center. Areas of the selected regions are as follows: ① 0.01 mm^2^ ② 2.08 mm^2^ ③ 2.08 mm^2^ ④ 2.09 mm^2^. The distance of each region to the centroid of the biofilm is also measured: ① 7.15 mm ② 6.77 mm ③ 6.12 mm ④ 6.56 mm. The scale bar measures 5 mm.

## 3. Results

### 3.1. Average spall size increases with increased laser fluence

The onset of film failure manifests differently depending on the type of film that is loaded. **Fig. 5** depicts the typical film failure progression for *S. mutans* biofilms, gold films, and sol-gel films. Failure for gold and sol-gel films begins by delamination from the substrate and “wrinkling” of the films. We hypothesize this “wrinkling” occurs at the onset of spallation for high-cohesive strength films. However, biofilms are a low-cohesive film, and the onset of failure is characterized by spallation of the bacteria without apparent “wrinkling” in the film. The failure region is more concentric for biofilms, and increases in size at increasing laser fluence. Failure for gold and sol-gel films evolves from wrinkling into film spallation at higher fluences, but the failure region is more irregular, due to the rupture of the films. As fluence continues to increase, all films exhibit spallation and an increase in spallation region size until the spall size reaches the diameter of the loading pulse. The maximum spallation region is equal to the loading pulse spot size. Thus, spall size is presented as a percent of possible spallation area based on the laser spot size.

Average and standard deviation of spall size is calculated for gold films, sol-gel films, and biofilms. The onset of spallation occurs at a different fluence for each film. At relatively low laser fluences, less than 40 mJ/mm^2^, biofilm rupture is infrequent and average spall size is markedly small. Spall size of biofilms increases monotonically with increasing laser fluence from 15.9 to 79.4 mJ/mm^2^. The gold and sol gel films were tested at different fluence ranges, 12.6 mJ/mm^2^ to 55.2 mJ/mm^2^ and 88.2 mJ/mm^2^ to 137.3 mJ/mm^2^, respectively. Because of different test fluence ranges, the graph shown in **Fig. 6** is normalized, for all films, to the initial fluence tested. It is important to note that the fluence values do not indicate which films exhibit greater adhesion. Interface stress is reliant on film-substrate properties including film thickness, density, and modulus of elasticity in addition to laser fluence. The gold films, sol-gel films, and biofilms exhibit a range of 0.73- 70.6%, 0-25.4%, and 0-45.1% of total spallation region failure over their respective fluence ranges. The normalized trend illuminates the similar failure modes found in the gold and sol-gel films. Both films initiate with film wrinkling and the measured spall size is very small. As fluence increases, wrinkling evolves into film rupture and spall size increases. However, film failure in biofilms is initiated directly by spallation and not a precursory “wrinkling” phenomenon.

**Fig. 5.**
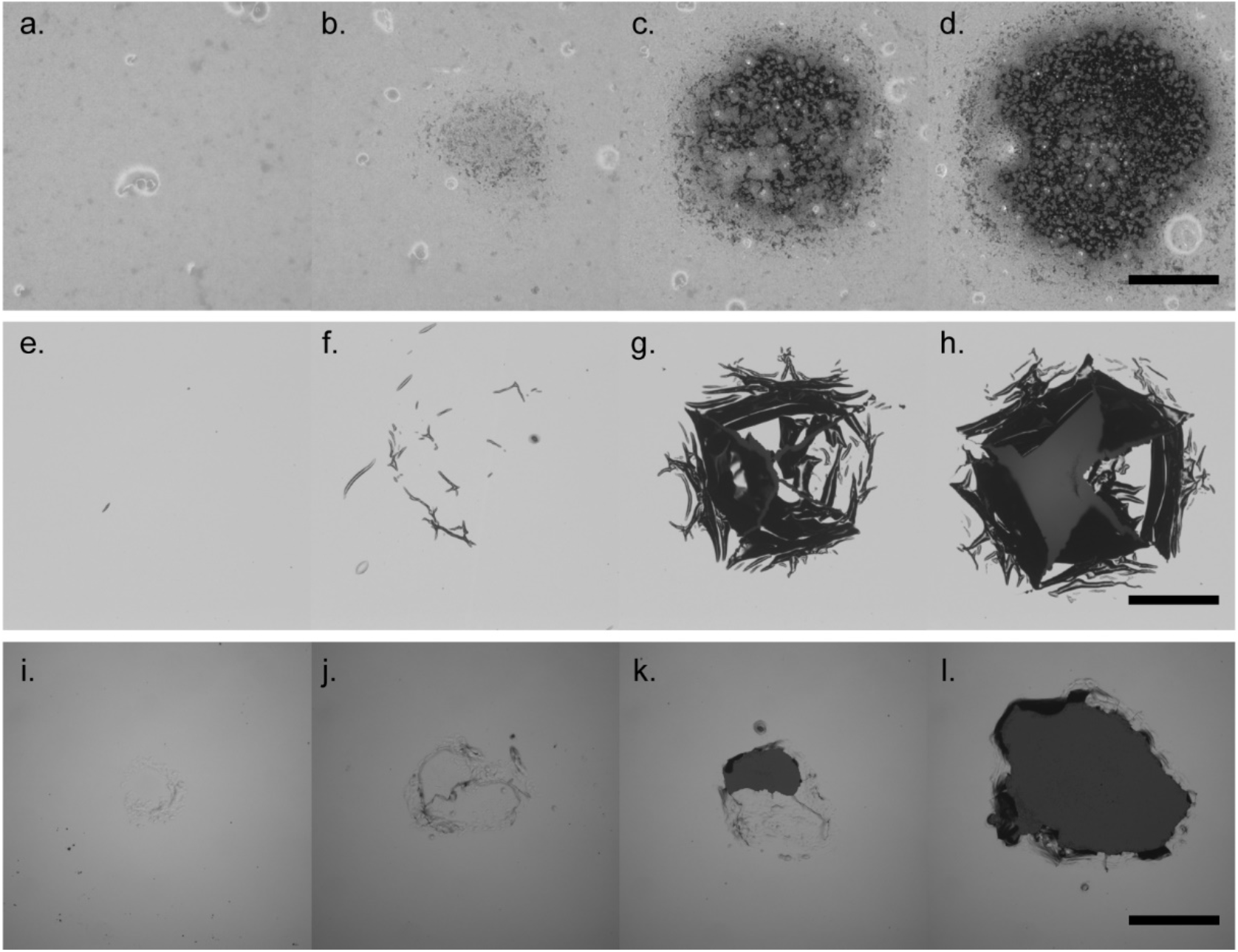
Optical images of failure progression in different films. (a-d) Depict failure of *S. mutans* biofilm cultured with 75 mM of sucrose, as fluence increases from 15.9 mJ/mm^2^ to 79.4 mJ/mm^2^, the scale bar represents 500 μm, (e-h) depicts failure progression for transfer printed gold film coated silicon substrate, from 9.99 mJ/mm^2^ to 24.1 mJ/mm^2^, the scale bar represents 500 μm, and (i-l) depicts the failure progression for sol-gel thin films coated in Pt/Ti superlayer from 88.2 mJ/mm^2^ to 137.3 mJ/mm^2^, the scale bar represents 250 μm.

**Fig. 6.**
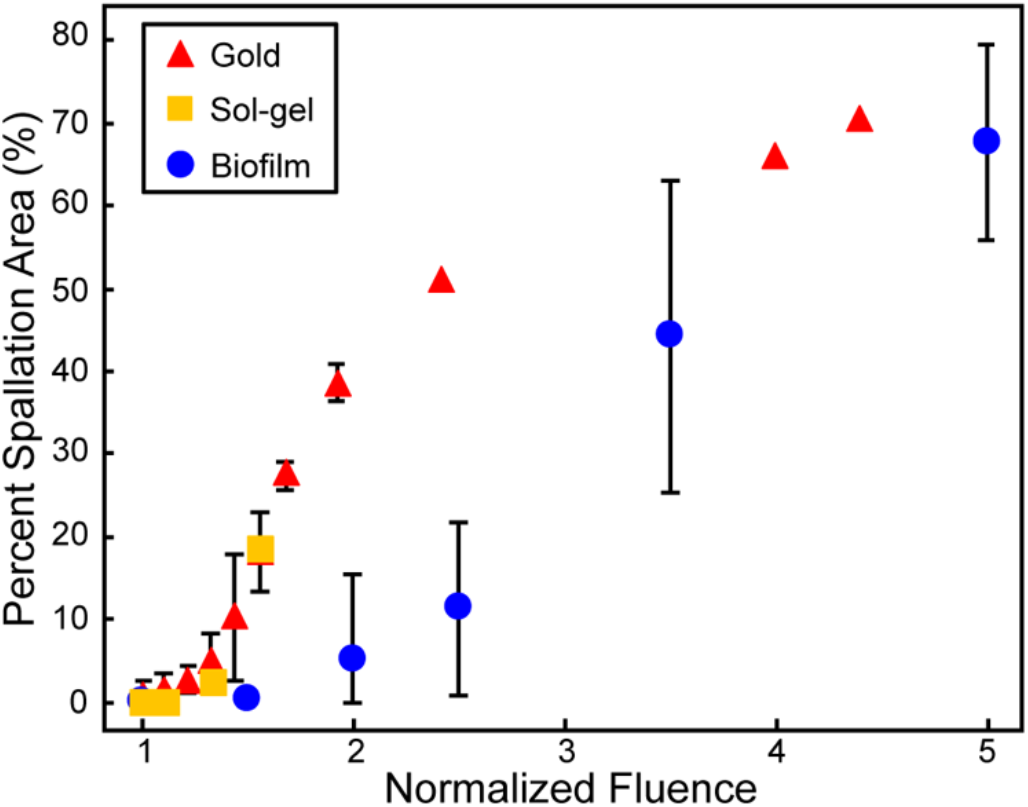
Scatter plots of percent spallation areas vs the normalized fluence applied to each film. The red triangles depict the statistics for the gold film (data from [25]), the yellow squares indicate the sol-gel film (data from [27]), and the blue circles indicate the failure for the biofilms tested. All of the films show a monotonically increasing relationship as fluence increases.

### 3.2 Spall area is independent of distance from centroid

During iso-fluence experiments, biofilms from 10 substrate assemblies are loaded 10-13 times each at either a laser fluence of 55.6 mJ/mm^2^ or 79.4 mJ/mm^2^. A total of 116 spallation regions are analyzed, which corresponds to 55 regions loaded at laser fluence 55.6 mJ/mm^2^ and 61 regions loaded at 79.4 mJ/mm^2^. A histogram of all spall size measurements for biofilms loaded at either fluence is shown in **Fig. 7**. The average and standard deviation of spall size for biofilms loaded at 55.6 mJ/mm^2^ is 1.19 ± 0.38 mm^2^ (31.2% of possible spall size) and for biofilms loaded at 79.4 mJ/mm^2^ is 1.69 ± 0.65 mm^2^ (44.4% of possible spall size). Spall size approximately follows a standard normal distribution for both fluence values. On average, an increase in laser fluence by 43% increases the spall size by approximately the same percentage, 42%. A two-sample Student’s t-test confirms independence of the two populations of fluence data with respect to spall size. The p-value is less than 0.0001, and thus is significant at that level.

**Fig. 7.**
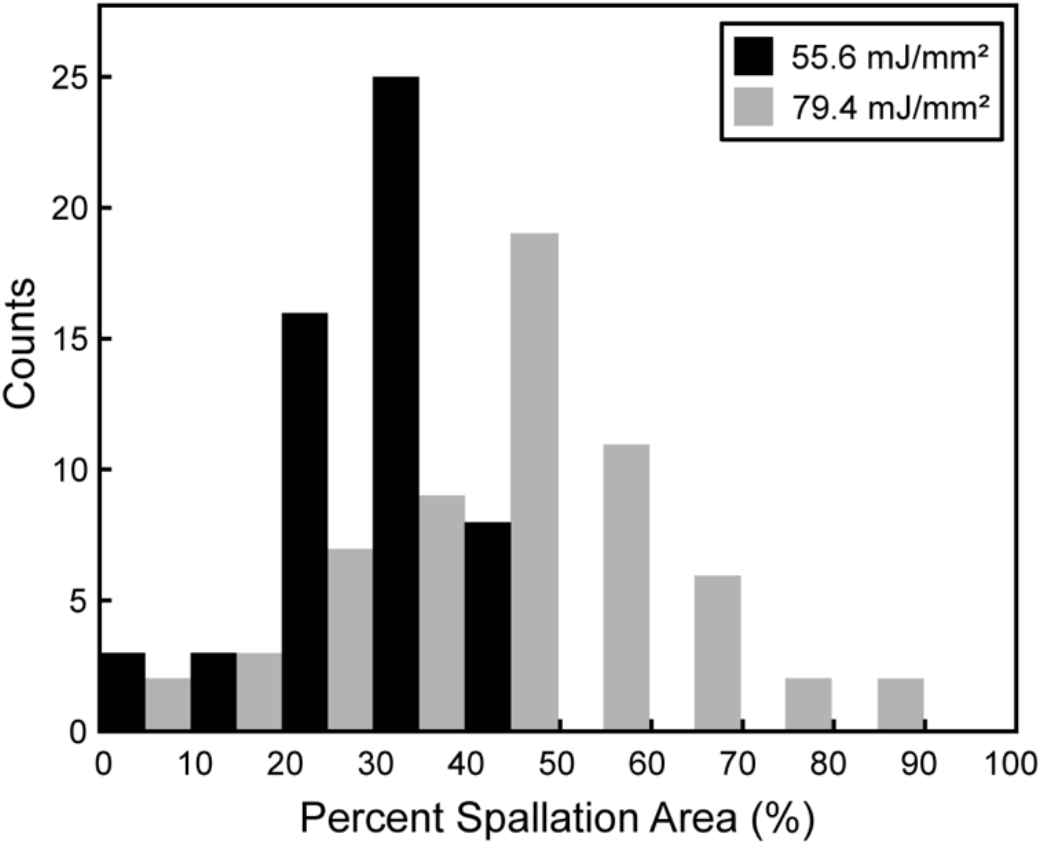
Histogram of percentage of possible spallation area for fluence of 55.6 mJ/mm^2^ (black bars) and 79.4 mJ/mm^2^ (gray bars).

On each biofilm, the centroid is located following image analysis protocol outlined in *Section 2.5*. The spall size in percent of possible spallation area is plotted with respect to distance from loading region to biofilm centroid in **Fig. 8**. The correlation between the two variables is measured using the Pearson correlation coefficient (PCC) method, which is advised for determining the relationship between two normally distributed variables [45]. The coefficients measured are *ρ* = −0.0994 and *ρ* = −0.0933 for biofilms loaded at 55.6 mJ/mm^2^ and at 79.4 mJ/mm^2^, respectively. The negative value for each coefficient indicates a slight negative relationship between the two variables. However, both PCC values fall within the 0.00 – 0.30 range, which Mukaka indicates as a negligible linear correlation [45]. In contrast with these values, the PCC between loading fluence and respective spall size is *ρ* = 0.6132, which indicates a moderate positive correlation. The PCC was calculated using the linear portion of biofilm data shown in **Fig. 6**. Data for regions loaded at a fluence below 30 mJ/mm^2^ were excluded to include only non-zero values for spall size. The number of loading locations at each loading distance is approximately constant except for within the first 2 mm from the biofilm centroid. There are fewer locations available for testing close to the center when compared to the number of locations available further from the center. It is a spatial limitation due to the 4 × 4 mm^2^ spacing of the loading locations.

**Fig. 8.**
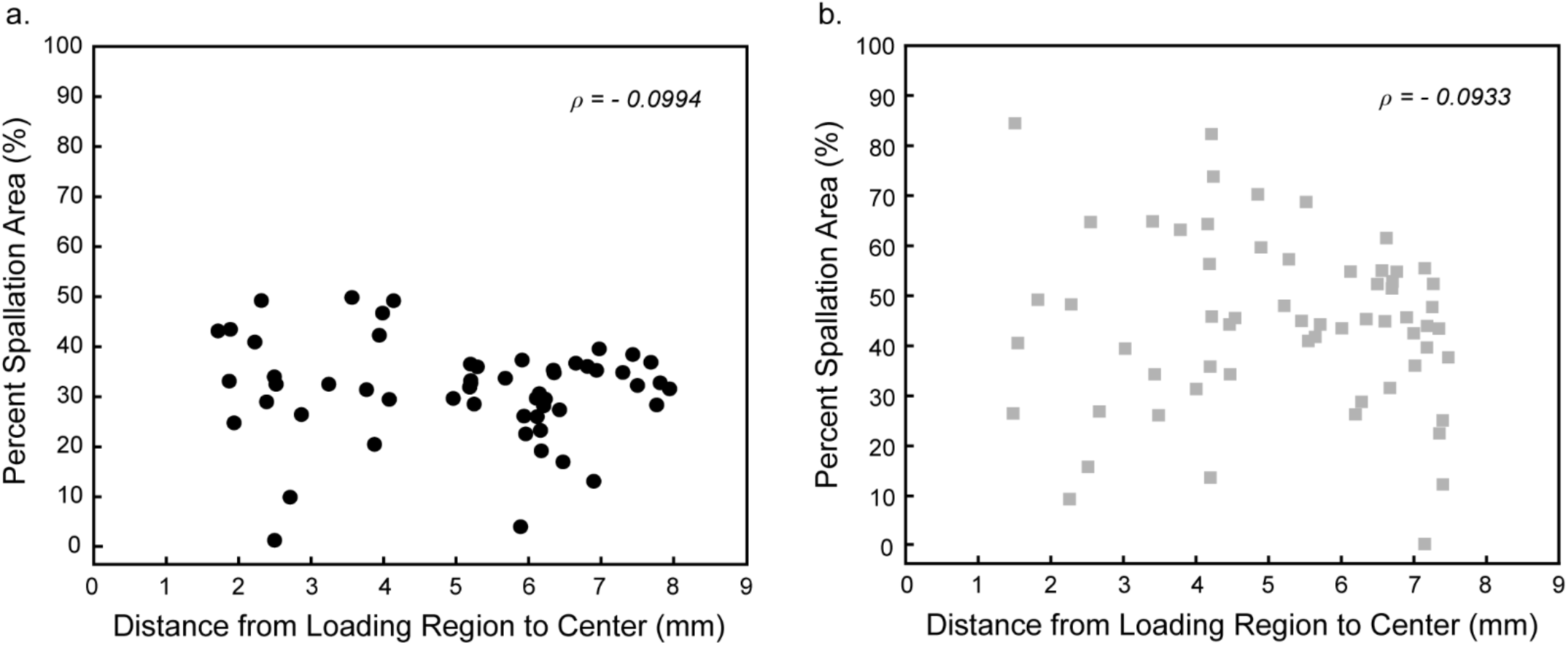
Scatter plots of percent spallation area with respect to their location on the sample for loading fluence of (a) 55.6 mJ/mm^2^ and (b) 79.4 mJ/mm^2^.

## 4. Discussion

After initial adhesion failure, an increase in laser fluence during variable fluence experiments is found to lead to a ubiquitous increase in spall size across three films: biofilms, gold films, and sol-gel films. The failure of these films presents differently, the gold and sol-gel films exhibit rapid increase in spall area, while biofilms require greater increases in fluence for similar increases in average failure area. Gold and sol-gel films exhibit film failure first by wrinkling before eventual rupture. Wrinkling of the films resulted in very small spall area values. For example, wrinkling in Fig. 5f and 5j indicate the film has delaminated from the substrate, but the detected spall size is negligible. In contrast, biofilm failure presents as spallation directly without film wrinkling (Fig. 5b), which lead to gradual increases in spall area with increases in laser fluence. Wrinkling at the onset of failure of metallic films loaded by laser-induced stress waves is an established phenomenon [33]. We hypothesize that the difference in failure mechanisms is associated with the cohesion of the film. High-cohesive films will tend to wrinkle at the onset of failure, while low-cohesive films will directly spall. At higher fluences, all three films experience spallation and display a convergence of film spallation regardless of film cohesion, which quantitatively confirms the trend identified in several publications [25–29].

Iso-fluence experiments are new in spallation literature. Our method to link spall area with distance to centroid is applied solely to our biofilms as this information was not available from the prior studies with gold and sol-gel films. The mean spall size for biofilms loaded at 55.6 mJ/mm^2^ is lower than that of biofilms loaded at 79.4 mJ/mm^2^. Additionally, spall area measurements at each loading fluence are normally distributed. The spread of spall sizes increased with increasing fluence, supporting the conclusion that fluence is directly correlated to the respective spall size. However, the statistical variability in spall area is quite large for biofilms. The range of areas loaded at 55.6 mJ/mm^2^ (1% to 50%) is encompassed by the range of areas loaded at 79.4 mJ/mm^2^ (0% to 85%). This large variation is likely due to the inherent variability in interface strength of the biofilms, which requires a higher number of tests to obtain statistical power. Through analysis of 116 images of loaded biofilm regions, we found no notable correlation between loading location and respective spall size for either loading fluence. The absolute value of PCC is less than 0.1 for both fluences, which indicates little to no relationship. Both loading fluences obtained similar PCC values, providing evidence that there is negligible correlation between the variables regardless of fluence. The slight negative correlation that appears to be present is attributed to biofilm heterogeneity.

## 5. Conclusions

In order to determine effects of biofilm membrane tension, we present a protocol that measures spall size with increasing laser fluence (variable fluence) and with respect to distance from the biofilm centroid (iso-fluence). *Streptococcus mutans* biofilms on titanium substrates are loaded via laser-induced stress waves through the laser spallation technique. A total of 185 biofilm loading locations are analyzed in this study. We demonstrate that biofilm spall size increases monotonically with laser fluence and apply our procedure to failure of non-biological films, which include gold films and sol-gel films. Different failure mechanisms are present among the three films, however, monotonic increases in spall area with fluence is a consistent trend across film types. Spall size measurements are dependent on the occurrence of film spallation. In films that present with “wrinkling” ahead of spallation, little to no measurable spall size is possible at lower loading fluences. This phenomenon deems adhesion strength measurements difficult through optical spall size measurements alone. However, because biofilms present with spallation at the onset of film rupture, the technique illustrated in this work is ideal for determining initial failure of biological films.

Because the laser spallation technique enables measurement of multiple locations across a biofilm, we can now determine the influence of membrane tension on adhesion measurements through iso-fluence experiments. We found no correlation between biofilm spall size and loading location, thus providing evidence that membrane tension does not play a dominant role in biofilm adhesion measurements. Furthermore, the methods described in this paper can be employed by other investigators to deconvolute location bias due to membrane tension within biological films or edge effects for synthetic films during laser spallation testing. We recommend our procedure as a straightforward method to determine membrane effects in the measurement of adhesion of biological films on substrate surfaces via the laser spallation technique. Preventing initial biofilm adhesion is a primary goal of new medical devices and having a consistent method for testing adhesion strength at the substrate-biofilm interface is extremely valuable.

## Acknowledgements

This work was supported by National Institutes of Health COBRE Phase III pilot funding under number 5P30GM110788-04. We thank the Center for Pharmaceutical Research and Innovation (CPRI) for use of bacterial culture equipment. CPRI is supported, in part, by the University of Kentucky College of Pharmacy and Center for Clinical and Translational Science (UL1TR001998). We would also like to thank Dr. Craig Miller from the University of Kentucky College of Dentistry for his guidance.

## Data Availability

The raw and processed data required to reproduce these findings are available to download from https://doi.org/10.18126/5LY6-ORUL via the Materials Data Facility [46, 47].

## References

1. Toda, Y., Moro, I., Koga, T., Asakawa, H., and Hamada, S., Ultrastructure of extracellular polysaccharides produced by serotype c Streptococcus mutans, Journal of Dental Research, 66 (8), 1364–1369 (1987).

2. Garrett, T. R., Bhakoo, M., and Zhang, Z., Bacterial adhesion and biofilms on surfaces, Progress in Natural Science, 18 (9), 1049–1056 (2008).

3. Klein, M. I., S Duarte, J Xiao, S Mitra, T H Foster, and H Koo, Structural and Molecular Basis of the Role of Starch and Sucrose in Streptococcus mutans Biofilm Development, Applied and Environmental Microbiology, 75 (3), 837–841 (2009).

4. Senadheera, M. D., Guggenheim, B., Spatafora, G. A., Huang, Y. C., Choi, J., Hung, D. C., Treglown, J. S., Goodman, S. D., Ellen, R. P., and Cvitkovitch, D. G., A VicRK Signal Transduction System in Streptococcus mutans Affects gtfBCD, gbpB, and ftf expression, biofilm formation, and genetic competence development, Journal of Bacteriology, 187 (12), 4064–4076 (2005).

5. Flemming, H. C. and Wingender, J., The biofilm matrix, Nature Reviews Microbiology, 8, 623 (2010).

6. Cross, S. E., Kreth, J., Zhu, L., Sullivan, R., Shi, W., Qi, F., and Gimzewski, J. K., Nanomechanical properties of glucans and associated cell-surface adhesion of Streptococcus mutans probed by atomic force microscopy under in situ conditions, Microbiology, 153 (9), 3124–3132 (2007).

7. Dunne, W. M., Bacterial Adhesion: Seen Any Good Biofilms Lately?, Clinical Microbiology Reviews, 15 (2), 155–166 (2002).

8. Waters, M. S., Kundu, S., Lin, N. J., and Lin-Gibson, S., Microstructure and Mechanical Properties of In Situ Streptococcus mutans Biofilms, ACS Applied Materials & Interfaces, 6 (1), 327–332 (2014).

9. Donlan, R. M., Biofilms and device-associated infections, Emerging Infectious Diseases, 7 (2), 277 (2001).

10. Zimmerli, W., Trampuz, A., and Ochsner, P. E., Prosthetic-joint infections, New England Journal of Medicine, 351 (16), 1645–1654 (2004).

11. Sridhar, S., Wang, F., Wilson, T. G., Palmer, K., Valderrama, P., and Rodrigues, D. C., The role of bacterial biofilm and mechanical forces in modulating dental implant failures, Journal of the Mechanical Behavior of Biomedical Materials, 92, 118–127 (2019).

12. Busscher, H., Rinastiti, M., Siswomihardjo, W., and Van der Mei, H., Biofilm formation on dental restorative and implant materials, Journal of Dental Research, 89 (7), 657–665 (2010).

13. Bürgers, R., Gerlach, T., Hahnel, S., Schwarz, F., Handel, G., and Gosau, M., In vivo and in vitro biofilm formation on two different titanium implant surfaces, Clinical Oral Implants Research, 21 (2), 156–164 (2010).

14. Hall-Stoodley, L., Costerton, J. W., and Stoodley, P., Bacterial biofilms: from the natural environment to infectious diseases, Nature Reviews Microbiology, 2 (2), 95 (2004).

15. Fux, C. A., Quigley, M., Worel, A., Post, C., Zimmerli, S., Ehrlich, G., and Veeh, R. H., Biofilm-related infections of cerebrospinal fluid shunts, Clinical Microbiology and Infection, 12 (4), 331–337 (2006).

16. Wu, H., Moser, C., Wang, H.-Z., Høiby, N., and Song, Z.-J., Strategies for combating bacterial biofilm infections, International Journal of Oral Science, 7 (1), 1 (2015).

17. Adair, C., Gorman, S., Feron, B., Byers, L., Jones, D., Goldsmith, C., Moore, J., Kerr, J., Curran, M., and Hogg, G., Implications of endotracheal tube biofilm for ventilator-associated pneumonia, Intensive Care Medicine, 25 (10), 1072–1076 (1999).

18. Cope, J. R., Collier, S. A., Nethercut, H., Jones, J. M., Yates, K., and Yoder, J. S., Risk behaviors for contact lens–related eye infections among adults and adolescents—United States, 2016, MMWR. Morbidity and Mortality Weekly Report, 66 (32), 841 (2017).

19. McLaughlin-Borlace, L., Stapleton, F., Matheson, M., and Dart, J., Bacterial biofilm on contact lenses and lens storage cases in wearers with microbial keratitis, Journal of Applied Microbiology, 84 (5), 827–838 (1998).

20. Weiss, L., The measurement of cell adhesion, Experimental Cell Research, 8, 141–153 (1961).

21. Stoodley, P., Cargo, R., Rupp, C. J., Wilson, S., and Klapper, I., Biofilm material properties as related to shear-induced deformation and detachment phenomena, Journal of Industrial Microbiology and Biotechnology, 29 (6), 361–367 (2002).

22. Gordon, V. D., Davis-Fields, M., Kovach, K., and Rodesney, C. A., Biofilms and mechanics: a review of experimental techniques and findings, Journal of Physics D: Applied Physics, 50 (22), 223002 (2017).

23. Katsikogianni, M. and Missirlis, Y., Concise review of mechanisms of bacterial adhesion to biomaterials and of techniques used in estimating bacteria-material interactions, Eur Cell Mater, 8 (3), 37–57 (2004).

24. Boyd, J., Korotkova, N., and Grady, M., Adhesion of Biofilms on Titanium Measured by Laser-Induced Spallation, Experimental Mechanics, 1–10 (2018).

25. Grady, M. E., Geubelle, P. H., Braun, P. V., and Sottos, N. R., Molecular tailoring of interfacial failure, Langmuir, 30 (37), 11096–11102 (2014).

26. Berfield, T. A., Kitey, R., and Kandula, S. S., Adhesion strength of lead zirconate titanate sol-gel thin films, Thin Solid Films, 598, 230–235 (2016).

27. Kandula, S. S. V., Hartfield, C. D., Geubelle, P. H., and Sottos, N. R., Adhesion strength measurement of polymer dielectric interfaces using laser spallation technique, Thin Solid Films, 516 (21), 7627–7635 (2008).

28. Navarro, A., Taylor, Z. D., Matolek, A. Z., Weltman, A., Ramaprasad, V., Huang, S., Beenhouwer, D. O., Haake, D. A., Gupta, V., and Grundfest, W. S. Bacterial biofilm disruption using laser-generated shockwaves. in SPIE BiOS. 2012. SPIE.

29. Gupta, V., Argon, A., Parks, D., and Cornie, J., Measurement of interface strength by a laser spallation technique, Journal of the Mechanics and Physics of Solids, 40 (1), 141–180 (1992).

30. Rickerby, D., A review of the methods for the measurement of coating-substrate adhesion, Surface and Coatings Technology, 36 (1–2), 541–557 (1988).

31. Chalker, P. R., Bull, S. J., and Rickerby, D. S., A review of the methods for the evaluation of coating-substrate adhesion, Materials Science and Engineering: A, 140, 583–592 (1991).

32. Stephens, A. W. and Vossen, J. L., Measurement of interfacial bond strength by laser spallation, Journal of Vacuum Science and Technology, 13, 38–39 (1976).

33. Wang, J., Weaver, R. L., and Sottos, N. R., A parametric study of laser induced thin film spallation, Experimental Mechanics, 42 (1), 74–83 (2002).

34. Grady, M. E., Geubelle, P. H., and Sottos, N. R., Interfacial adhesion of photodefinable polyimide films on passivated silicon, Thin Solid Films, 552, 116–123 (2014).

35. Vogel, A. and Venugopalan, V., Mechanisms of pulsed laser ablation of biological tissues, Chemical Reviews, 103 (2), 577–644 (2003).

36. Hu, L., Zhang, X., Miller, P., Ozkan, M., Ozkan, C., and Wang, J., Cell adhesion measurement by laser-induced stress waves, Journal of Applied Physics, 100 (8), 084701 (2006).

37. Hagerman, E., Shim, J., Gupta, V., and Wu, B., Evaluation of laser spallation as a technique for measurement of cell adhesion strength, Journal of Biomedical Materials Research Part A, 82 (4), 852–860 (2007).

38. Stoodley, P., Lewandowski, Z., Boyle, J. D., and Lappin‐Scott, H. M., Structural deformation of bacterial biofilms caused by short‐term fluctuations in fluid shear: An in situ investigation of biofilm rheology, Biotechnology and Bioengineering, 65 (1), 83–92 (1999).

39. Kumar, P. S., Mason, M. R., Brooker, M. R., and O’Brien, K., Pyrosequencing reveals unique microbial signatures associated with healthy and failing dental implants, Journal of Clinical Periodontology, 39 (5), 425–433 (2012).

40. Elias, C., Lima, J., Valiev, R., and Meyers, M., Biomedical applications of titanium and its alloys, JOM, 60 (3), 46–49 (2008).

41. Norowski Jr, P. A. and Bumgardner, J. D., Biomaterial and antibiotic strategies for peri‐ implantitis: A review, Journal of Biomedical Materials Research Part B: Applied Biomaterials, 88 (2), 530–543 (2009).

42. Ferraris, S. and Spriano, S., Antibacterial titanium surfaces for medical implants, Materials Science and Engineering: C, 61, 965–978 (2016).

43. Besinis, A., Hadi, S. D., Le, H., Tredwin, C., and Handy, R., Antibacterial activity and biofilm inhibition by surface modified titanium alloy medical implants following application of silver, titanium dioxide and hydroxyapatite nanocoatings, Nanotoxicology, 11 (3), 327–338 (2017).

44. Schneider, C. A., Rasband, W. S., and Eliceiri, K. W., NIH Image to ImageJ: 25 years of image analysis, Nature Methods, 9, 671 (2012).

45. Mukaka, M. M., A guide to appropriate use of correlation coefficient in medical research, Malawi Medical Journal, 24 (3), 69–71 (2012).

46. Blaiszik, B., Chard, K., Pruyne, J., Ananthakrishnan, R., Tuecke, S., and Foster, I., The Materials Data Facility: Data services to advance materials science research, JOM, 68 (8), 2045–2052 (2016).

47. Blaiszik, B., Ward, L., Schwarting, M., Gaff, J., Chard, R., Pike, D., Chard, K., and Foster, I., A data ecosystem to support machine learning in materials science, MRS Communications, 1–9 (2019).

